# The trifecta of disease avoidance, silique shattering resistance and flowering period elongation achieved by the *BnaIDA* editing in *Brassica napus*

**DOI:** 10.1101/2022.06.09.493885

**Authors:** Rui Geng, Yue Shan, Lei Li, Chun-Lin Shi, Jin Wang, Wei Zhang, Rehman Sarwar, Yi-Xuan Xue, Yu-Long Li, Ke-Ming Zhu, Zheng Wang, Li-Zhang Xu, Reidunn B. Aalen, Xiao-Li Tan

## Abstract

Rapeseed (*Brassica napus*) oil is a main vegetable oil source in the world. The devastating disease of stem rot caused by the necrotrophic fungus *Sclerotinia sclerotiorum* and pod shattering led to a great yield loss in *Brassica napus. S*.*sclerotiorum* infects the rapeseed by the detached floral petals, in which the released ascospores land and germinate as mycelium, then the petals fall on the leaves at lower part of the rapeseed and heavily attacks the leaves and stems. The prevention of petal-shedding is a promising approach to avoid the stem rot damage, moreover, longer period of flowering time will bring rapeseed flower tourism a huge economic benefit. Notably, IDA (INFLORESCENCE DEFICIENT IN ABSCISSION) and IDA-LIKE(IDL) protein control floral organ abscission in *Arabidopsis thaliana*. In our study, the precisely editing of two IDA homologues genes using CRISPR/Cas9 system in *Brassica napus* caused the petal attaching to the flower till pod mature and enhancing the silique dehiscence resistance. Incubating the *S.sclerotiorum* to petal showed the edited rapeseed avoiding the infection of *S.sclerotiorum* RNA-Seq analysis demonstrated that in the editted plant, the genes involed in IDA pathway were regulated, while other genes keep unaltered. Investigation of agronomic traits showed that no positive the agronimic traits was introduced in editted plant. Our study demonstrated that mutation of two BnaIDAs creating a promising germplasm for disease avoidance, siliques shattering resistance and flowering period elongation which will contribute great to rapeseed industry.

## Introduction

Rapeseed (*Brassica napus*) oil is a main vegetable oil source in the world. The devastating disease of stem rot caused by the necrotrophic fungus *Sclerotinia sclerotiorum* and pod shattering lead to a great yield loss in *Brassica napus*(X. Zhang et al., 2021). *S*.*sclerotiorum* infects the rapeseed by the detached floral petals. The released ascospores land on the petal and germinate as mycelium, then the petals fall on the leaves at lower part of the rapeseed and heavily attacks the leaves and stems (Bolton et al., 2006; L. N. Ding et al., 2021). Blocking the petal mediated infection is a promising approach to avoid the stem rot damage. For instance, apetalous rapeseed decreased the infections (D. I. Jamaux, and Spire, D., 1999; K. Yu et al., 2018). The prevention of floral shedding may reduce the further infection, moreover, it will bring rapeseed flower tourism a huge economic benefit due to longer period of flowering time. The easy silique dehiscence caused the yield loss up to 50% during the mechanized harvesting of *Brassica napus* (Li et al., 2021). IDA (INFLORESCENCE DEFICIENT IN ABSCISSION) and IDA-LIKE(IDL) protein control floral organ abscission and silique dehiscence in *Arabidopsis* (Butenko et al., 2003; Santiago et al., 2016). The story of IDA in Arbidopsis shed a light on rapeseed to generate a germplasm with attached petals and shattering proof siliques.

The overall detachment of plant organs is defined as abscission. There are many factors affecting the beginning of abscission, the most common is hormones, such as the abscission process of floral organs can be regulated by ethylene. Studies in *Arabidopsis* showed that ethylene insensitive mutants (*ein1* and *ein2*) significantly delayed the abscission of floral organs (Guzman et al., 1990). And auxin also plays a key role in organ abscission, it is found that auxin may regulate abscission through changes in the flow distribution of abscission zone in tomato (Dong et al., 2021). In addition to plant hormones as signal molecules to regulate plant organ abscission, a small peptide is required for abscission is IDA (INFLORESCENCE DEFICIENT IN ABSCISSION) and IDA-LIKE(IDL) protein families composed of no more than 100 amino acids are also used as signal molecules to participate in the regulation of floral organ abscission (Butenko et al., 2003; Santiago et al., 2016). In IDA-HAE/HSL2 pathway, the polypeptide ligand IDA binds to the LRR protein kinase HAE/HSL2 co-receptor on the cell membrane, and then phosphorylates after being recognized by

SOMATIC EMBRYOGENESIS RECEPTOR KINASES(SERKs), so as to activate the downstream MITOGEN-ACTIVATED PROTEIN KINASE (MAPK) protein kinase cascade. Then, the activities of transcription factors such as *BREVIPEDICELLUS (BP)/KNOTTED-LIKE FROM ARABIDOPSIS THALIANA1 (KNAT1)* and *KNAT2/6* are regulated, and finally control the hydrolase activity in the cell wall and regulate floral organ abscission (Butenko et al., 2003; Cho et al., 2008; Lewis et al., 2010; Meng et al., 2016; Santiago et al., 2016; Shi et al., 2011; Stenvik et al., 2008). In addition, IDA/IDL also participates in the process of lateral root emergence and root abscission, and is closely related to the maturation of the disjunction with lateral root primordia (LRP) and lateral root cap(LRC) cells(Shi et al., 2019). Studies in *Nicotiana benthamiana* showed that IDA was also involved in water stress and stem growth and development (Ventimilla et al., 2020). It seems that IDA/IDL as a conserved signal peptide is involved in cell activities in the abscission zone of plant life.

Rapeseed, which can survive in both high and low latitudes, is one of the most widely cultivated edible oil crops in the world. In particular, *Brassica napus* (AACC, 2n=38) is an allotetraploid hybrid evolved from the hybridization of *Brassica rapa* (AA, 2n=20) and *Brassica oleracea* (CC, 2n=18), belongs to Cruciferae with *Arabidopsis*(Chalhoub et al., 2014). However, the complex genome structure makes it difficult to obtain effective and specific genotype or phenotype mutations in *B*.*napus*, Unlike *Arabidopsis*, due to the existence of homologous genes, it is necessary to mutate multiple genes to find obvious phenotypes in *B*.*napus*. Nowadays, CRISPR/Cas9, a powerful efficient tool, has been used in various crops in modern agriculture and applied to various problems such as yield, plant type and disease. Previously, three genes (*SlER, SP5G* and *SP*) controlling stem length and flowering and fruiting time of tomato were modified by gene editing tools, and a new type of “urban tomato” was created by integrating excellent traits (Kwon et al., 2020). Genome editing of the Glucan, Water-Dikinase 1(GWD1) in rice showed that the gene has good application potential to improve the yield and quality of rice at the same time(Wang et al., 2021). Gene editing is also widely used in *B*.*napus*, knockout of *BnaLPAT2/5* gene can change the oil body size and fatty acid content, then genome editing of *BnaQCR8* can improve the resistance to two fungi and will not affect other excellent agronomic traits(K. Zhang et al., 2019; X. Zhang et al., 2021).

Premature abscission of floral organs or silique is a major constraint on crop production, especially in *Brassica napus*, which cause shorten the ornamental flowering period, and the yield loss can be up to 50% in mechanized harvesting (Li et al., 2021). Furthermore, the spread of stem rot in *B*.*napus* is caused by *S*.*sclerotiorum*, which will cause a loss of 8.4-billion-yuan RMB only in China every year (X. Zhang et al., 2021). Hence, the creation of new germplasm with multiple excellent traits is necessary for the development of rapeseed industry in modern society.

In order to study the role of secretory small peptide IDA in *Brassica napus* and explore its application value, we used CRISPR/Cas9 technology to edit the two *BnaIDA* genes, which most similar to *Arabidopsis thaliana*, resulting in the petal attaching to the flower till pod mature and the silique dehiscence resistance.

Removing exogenous Cas9 by selfed crossing the mutants, and designed primers for rapid identification, which proved that the mutation caused by gene editing can be inherited stably. We were surprised to find that *Bnaida* also has high silique dehiscence and displayed prevent disease phenotype to *S. sclerotiorum*, which has high application value. In this study, we have precisely edited the *BnaIDA-A07/C06* gene through CRISPR/Cas9 technology, and gathered three excellent traits on one plant through one transformation event.

## Results

### Knockout of *IDA* genes in *Brassica napus*

The gene and protein sequences of IDA in Arabidopsis (Shi et al., 2011) were obtained and analyzed by *B*.*napus* database(https://www.genoscope.cns.fr/brassicanapus/). The five IDA homologous genes in *Brassica napus* were obtained. We named *BnaIDA-A07*(BnaA07g27400D), *BnaIDA-C06*(BnaC06g29530D), *BnaIDA-C02*(BnaC02g18450D), *BnaIDA-C04*(BnaC04g26010D) and *BnaIDA-A02*(BnaA02g13980D). The identities with AtIDA were 88.3%, 89.2%, 85.3%, 86.8% and 87.0% respectively. *BnaIDA-A07* and *BnaIDA-C06* had the highest similarity with *AtIDA* (Figure 1a), and the expressions of two genes are mainly in flowers and mature siliques (Figure 1b). *B*.*napus* transcriptome database was mined (https://brassica.biodb.org/) and results showed that *BnaIDA-A07* and *BnaIDA-C06* were specifically expressed in the floral organs, and the expression patterns of the other three homologous genes were different. The above results indicate that *BnaIDA-A07* and *BnaIDA-C06* were involved in the process of floral organs non-abscission *B*.*napus* (Figure S1).

**Figure 1.**
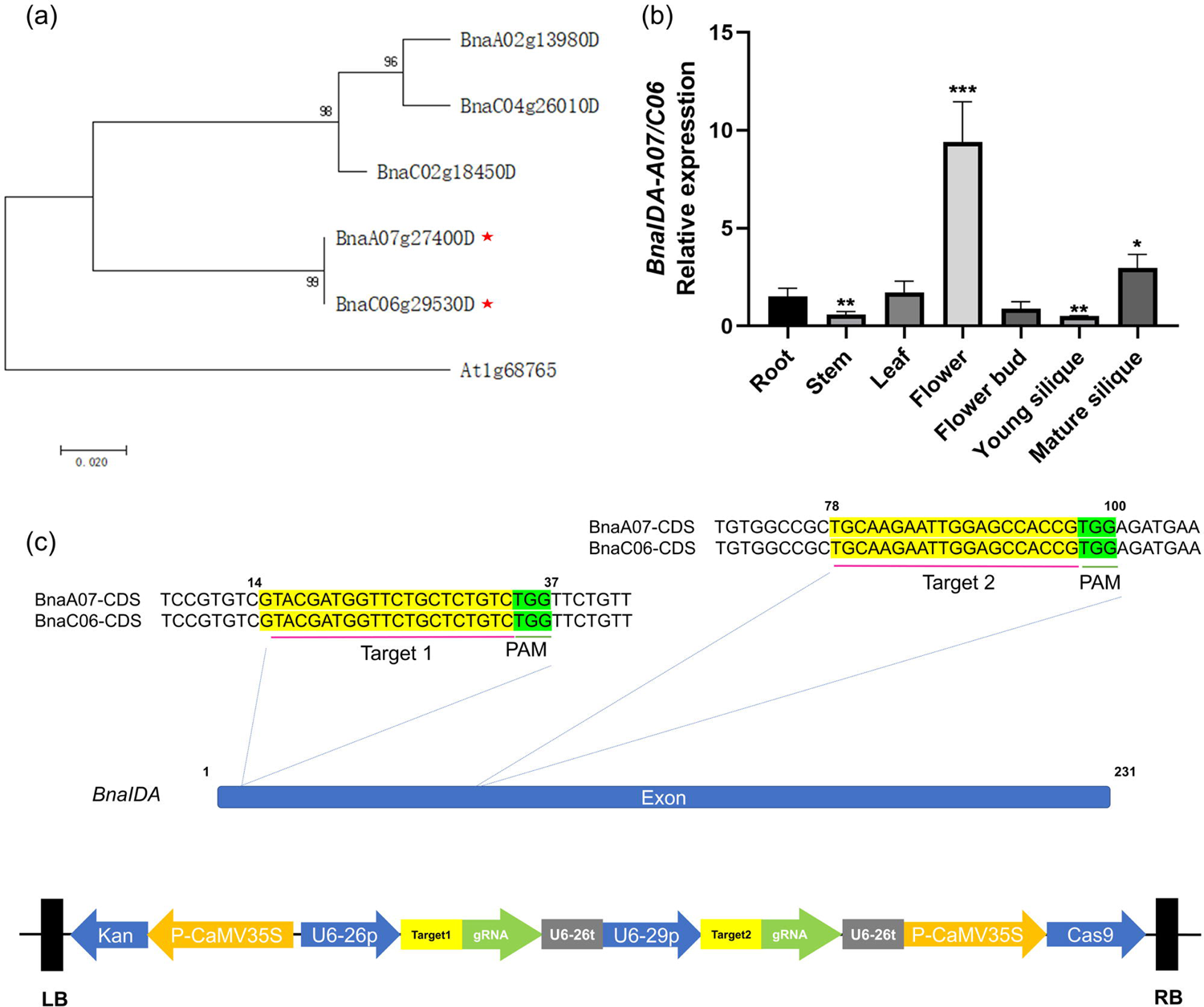
Phylogenetic relationships of *BnaIDA* homologous genes and strategy of gene editing. (a) The phylogenetic tree was made with the neighbor-joining algorithm. The Bootstrap value was used to check the branching reliability of phylogenetic tree. It could be seen that BnaA07g27400D and BnaC06g29530D were the most homologous with At1g68765. (b) Quantitative RT-PCR analysis of the expression of *BnaIDA-A07/C06* genes in various tissues of WT. BnaIDA-A07/C06 sequence has high homology and cannot be distinguished by the same primer. The significances of the gene expression differences are indicated (*** Student’s t-test, P > 0.001; ** Student’s t-test, P > 0.01; * Student’s t-test, P > 0.05; n=3, bars = SD). (c) Diagram of the targeted region of *BnaIDA-A07/C06* and the CRISPR-Cas9 vector. Targeted sites in the conserved CDS region of *BnaIDA-A07/C06* were presented. The U6-26 promoter was used to drive the targeting-sequences-fused sgRNA expression. Kanamycin resistance was used for the selective marker.

To specifically knock out the *BnaIDA-A07* and *BnaIDA-C06* gene simultaneously, two targets were deliberately selected in CDS region to ensure effective editing, which was likely to cause frameshift mutation. Two sequences targeting on the conserved region were constructed based on the CRISPR/Cas9 multiplex editing system (Y. Ding et al., 2016). SgRNAs scaffold were driven by Ubiquitin 6-26/29 promoter, while the Cas9 protein and Kanamycin resistance gene was under the control of *Cauliflower mosaic virus* 35S (Figure 1c).

Four positive complete plantlets identified by PCR were obtained after bud differentiation and rooting culture. PCR amplification was performed using the Ubiquitin 6-26/29 promoter and Cas9 sequence primers (Figure S2). Four T0 transgenic lines were self-pollinated to produce T1 progeny lines. Furthermore, for check the editing efficiency, *BnaIDA-A07* and *BnaIDA-C06* were effectively edited in both T0 and T1 lines confirmed by Sanger sequencing. The editing types were single or multi bases insertions or deletions (Figure 2a,2b). Notably, at target 1 site, the frequency of one T base insertion was 48%; at target 2 site, the frequency of one base deletion and one A base insertion was 38% and 26%, respectively (Figure 2c). Seven plants showed that editing had occurred in genome (Figure 2b, S2) without Cas9 and U6 promotor identified with PCR. The editing positions in the genome are accurately located near the PAM sequence. These results imply that simultaneous knockout of multiple genes in *B*.*napus* can be completed, and the exogenous U6 promoter and Cas9 protein can be isolated from the offspring through selfing, which indicate that gene editing can be used for rapid breeding.

**Figure 2.**
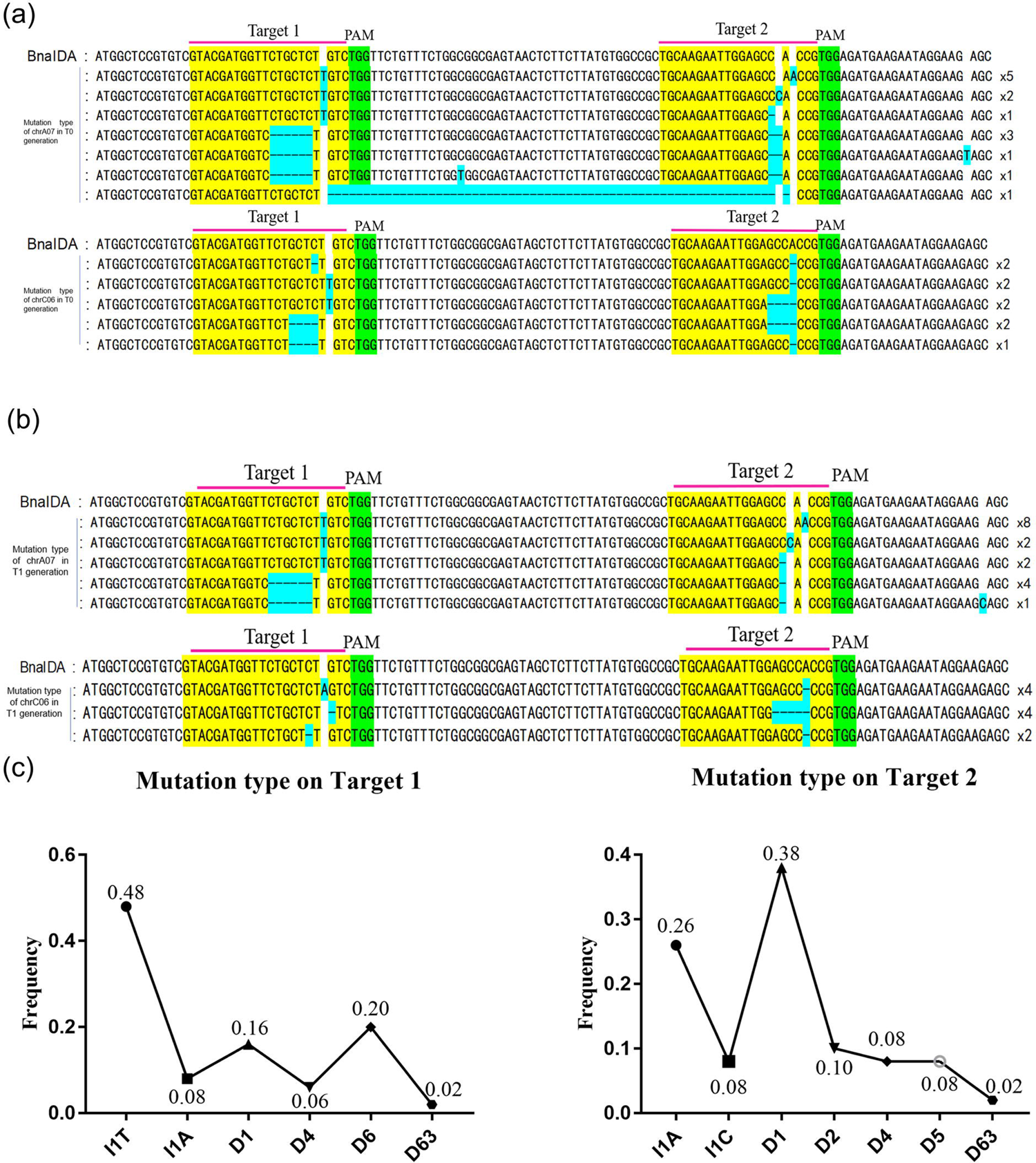
Genotype of *BnaIDA*-edited T0 and T1 mutants of rapeseed. Four *BnaIDA*-edited T0 plants(a) and seven *BnaIDA*-edited T1 plants(b) showing mutation sites at two homologous copies of *BnaIDA* (*BnaIDA-A07/C06*). Target were coloured yellow. Light blue represented the edited base. PAMs were coloured green. “-” indicated nucleotide deletions in the target sequence. The last number in each sequence represents the number of repetitions. (c) Mutation types and frequency at the sgRNA target sites in T0 and T1 mutants. The X-axis: I# and D# were the numbers of base pairs inserted and deleted at the sgRNA target sites.

In addition, the insertion of 1 bp(T) at Target 1, directly lead to the frameshift mutation of *BnaIDA-A07/C06* and the premature termination of the translation process, resulting in the inactivation of BnaIDA protein (Figure 3a). To easily distinguish WT and *Bnaida* mutant alleles and quickly identify the mutants with editing at two targets, allele specific PCR markers were developed. As previously mentioned, 1 bp(T) inserted in Target 1 and 1 bp(A) inserted in Target 2, The terminal base of primer 3’ had a great influence on the DNA synthesis efficiency of Taq enzyme (Wu et al., 2020). At target 1, we added T at the end of 3’ in forward primer, and at target 2 added A at the end of 3’ in reverse primer, named T2-F/A2-R(Figure 3b). Allele specific PCR primers were designed to identify WT and *Bnaida* mutant alleles. We propagated the above-mentioned seven T1 plants to obtain T2 plants. The homozygous editing events in the genome were confirmed by PCR and the phenotype showing no petals abscission. (Figure 3c and 4a). Thus, WT and homozygous mutants can be easily distinguished based on genotype identification.

**Figure 3.**
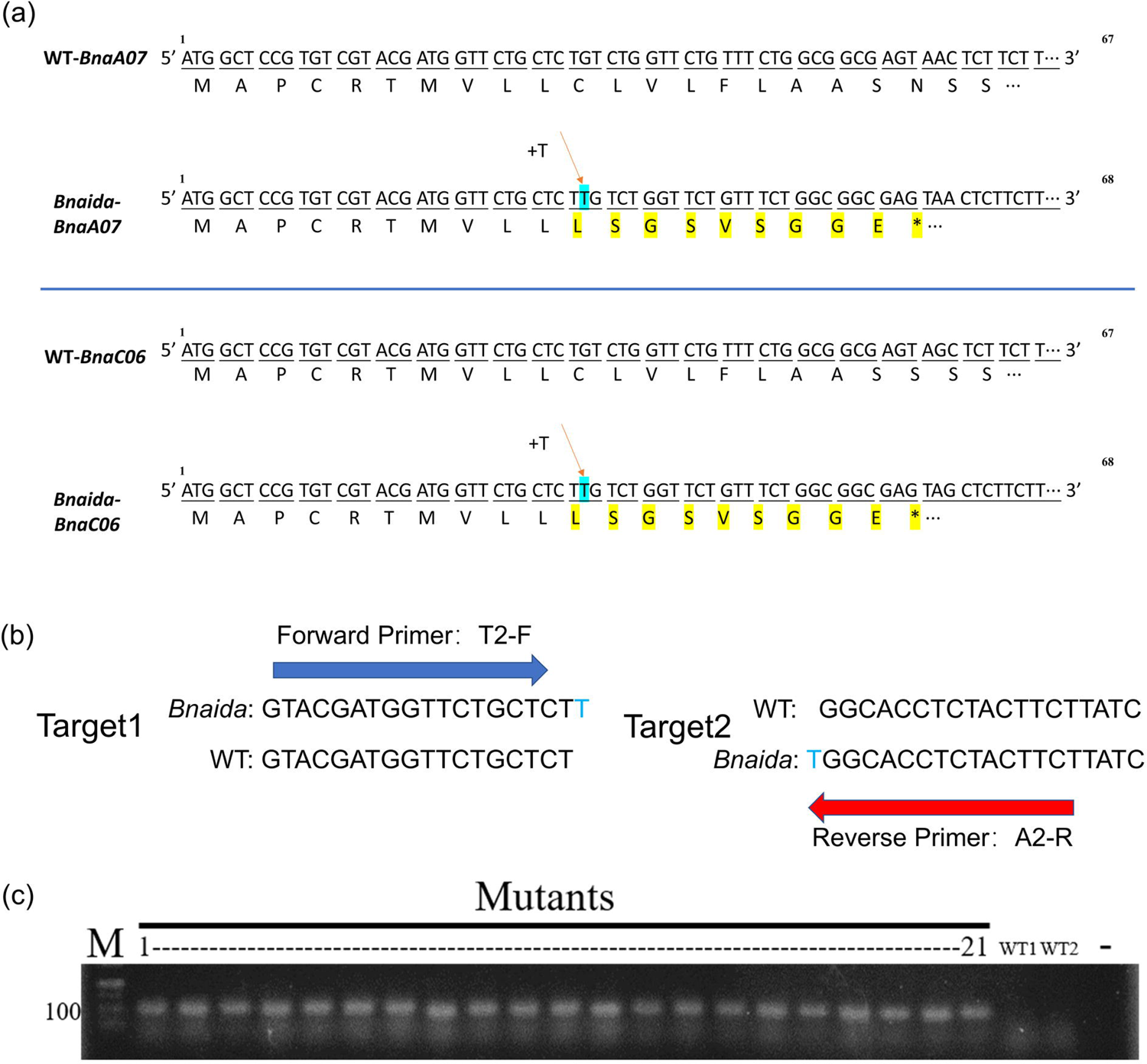
Rapid identification based on genotype. (a) Schematic diagram of frame shift mutation. Light blue represented the edited base, amino acid changes due to frameshift mutations were highlighted in yellow. (b, c) Design of primers for rapid identification of CRISPR edited sequences. One single base “T” in light blue was added at the 3’ end of both the forward and reverse primers. The CRISPR edited T2 lines were screened by showing PCR products while no bands were amplified in WT. “-”: ddH2O.

**Figure 4.**
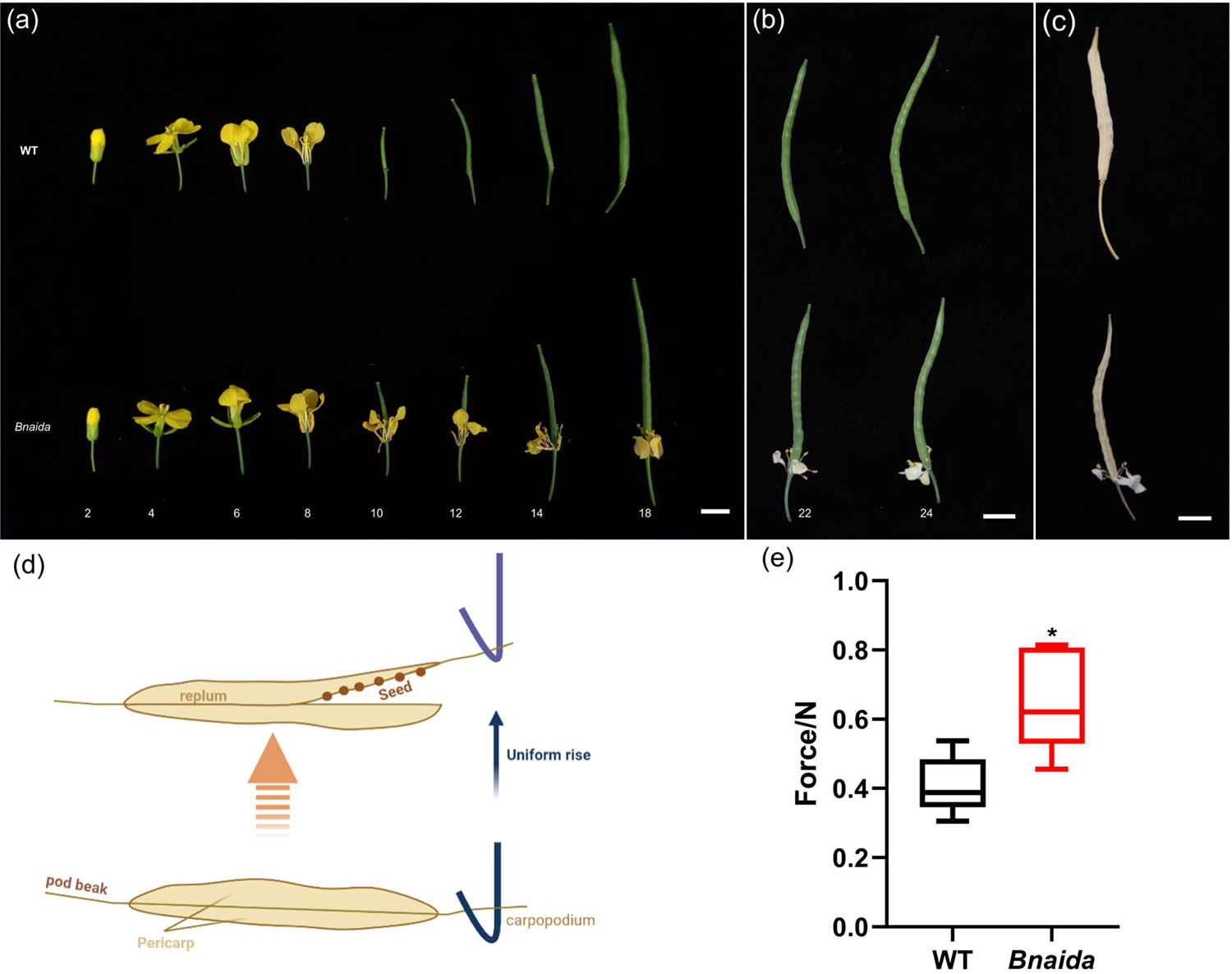
Characterization of *Bnaida* knock-out mutants. (a, b and c) WT plants occurred floral abscission shortly after anthesis, which defines position 1/2 when counting florals along the primary inflorescence. After the stage of position 10, only *Bnaida* retained floral organs indefinitely, even when the pod was fully mature and dry, bars = 1cm. (d, e) Schematic diagram of silique dehiscence-resistance force test. The pods were fixed on the plate, and the hook would pull the carpopodium during the uniform upward movement, resulting in the cracking of the pod. The instantaneous cracking force was measured by the physical property instrument. Forces of *BnaIDA* mutants were compared with WT pods, respectively. (n = 5, bars = SD; * Student’s t-test, P > 0.05).

### The flowering period of *Bnaida* was prolonged and enhances the resistance to silique dehiscence in *B*.*napus*

We counted the first flower at the top of the inflorescence as position 1 (Butenko et al., 2003). At the stage of position 1, the first flower can be visible at the top of the inflorescence with the yellow petals starting to appear from the sepals (Butenko et al., 2003). At the positions 2, 4, 6 and 8, the stigma was pollinated and began to expand. WT floral organs abscised very soon, showing completely no petals at the stage of position 10 (Figure 4a). In contrast, the floral organs of *Bnaida-A07/C06* mutants remained attached during the whole flowering to maturation stages (Figure 4b), even when the silique completely dried up, the petal still attached to the silique and the color of petals changed to white (Figure 4c). The IDA-HAE/HSL2 floral abscission signaling pathway leads to organ separation by affecting the cell activity of AZ (Abscission Zone). Dehiscence of silique was also a process of organ separation, previous studies had shown that organ detachment was associated with the expansion of AZ cells (Stenvik et al., 2006). In order to test the effect of *Bnaida* on silique dehiscence-resistance, the silique from WT and mutant plants at turning to light yellow stage were selected (Li et al., 2021), and the force required for silique dehiscence of *B*.*napus* was measured by texture analyzer (Figure 4d). The maximum tensile strength of WT was 0.3-0.5N and that of mutant was 0.6-0.8N (Figure 4d, S3), indicating that silique dehiscence-resistance was enhanced significantly due to the knockout of *BnaIDA-A07/C06*. The results showed that opening the siliques of *Bnaida* required more force, indicating that its siliques were shattering resistant.

### *BnaIDA-A07/C06*-edited mutants displayed prevent disease phenotype to *S. sclerotiorum*

*S*.*sclerotiorum* infection of rapeseed was spread by the fallen petals (Bolton et al., 2006; L. N. Ding et al., 2021). Since the petals of the mutants closely attached to the flower and did not fall onto the lower leaves. The petals of mutants and WT were inoculated with *S*.*sclerotiorum* to investigate the influence of floral abscission on stem rot disease development. The inoculated petals of WT were removed and put onto the leaves 1day post-inoculation (dpi) and the infected petals of mutants remained on the plants (Figure 5). As shown in Figure 5 almost all leaves of WT plants developed soft-rotting necrosis while no obvious disease development was observed either in petals or leaves of the mutants 4 dpi. Therefore, non-abscission of petals blocked the way to further spread of *S*.*sclerotiorum* and thus prevented the prevalence of the disease.

**Figure 5.**
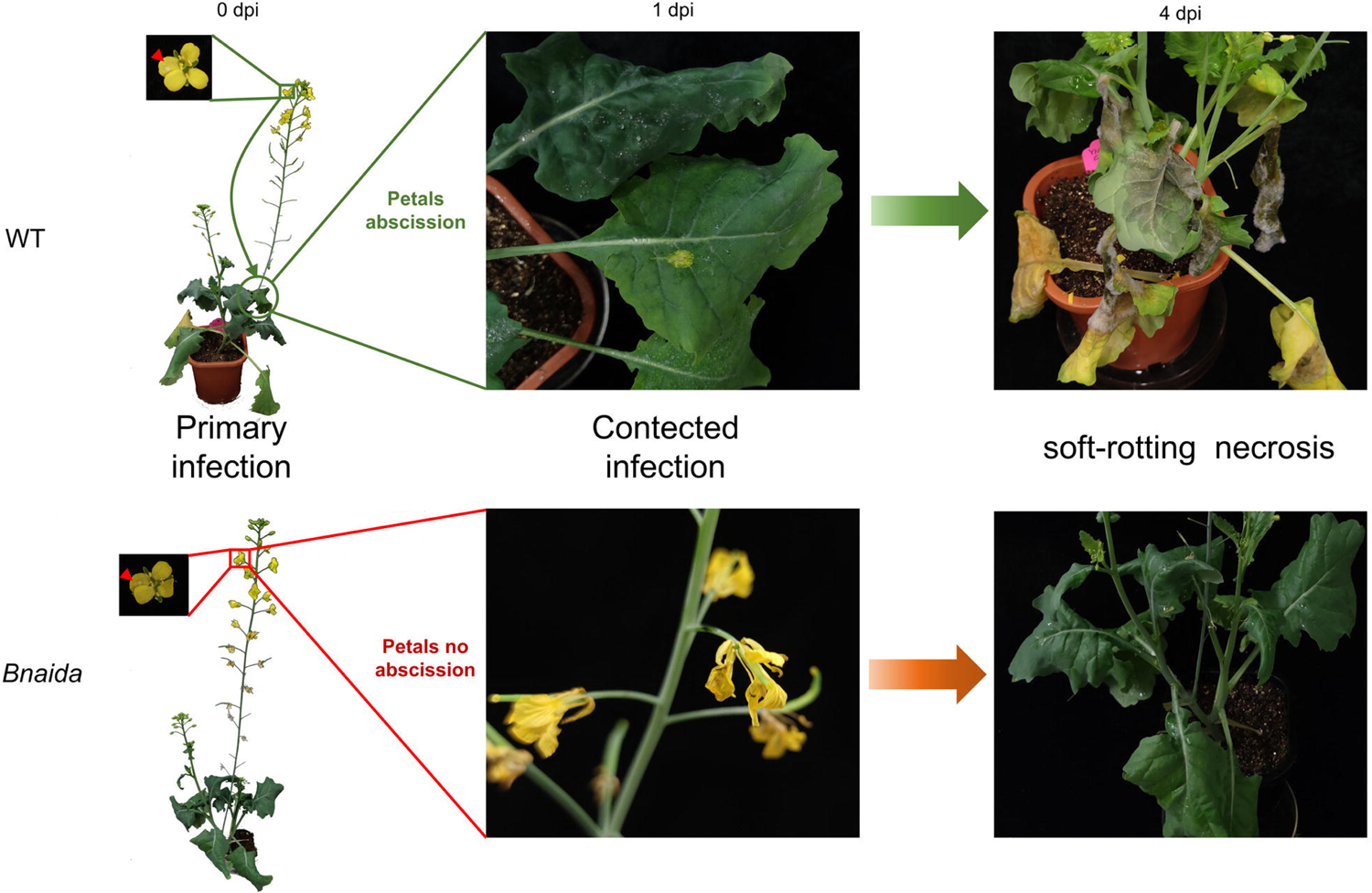
*Bnaida* mutant lines prevented disease spread of *S*.*sclerotiorum. S*.*s* avoidance of mutants with non-shedding floral organs. Disease responses of inoculated plants 0,1,4 dpi; The red arrows show the location of *Ss*, bars = 5 cm.

### Gene editing with *BnaIDA-A07/C06* affected the isolation of AZ cells

To further investigate whether the absence of IDA protein influences AZ (ABSCISSON ZONE cells), the abscission zone of WT and *Bnaida* mutants was observed under scanning electron microscopy at different stages of inflorescence development. Different from the gradual expansion and roundness of cells in the AZ of *Arabidopsis* (Liu et al., 2013). Before position 7/8, the floral organs of wild-type and *Bnaida* did not fall off, while after position 9/10, only the floral organs of *Bnaida* remained attached (Patterson et al., 2004). In general, different from the gradual expansion and roundness of cells in the abscission zone of Arabidopsis (Liu et al., 2013), the surface of WT residual AZ cells gradually became flat as a whole during the process of flower development and no petal, stamen or sepal abscission zone cells remained and could be transdifferentiated into epidermal cells in *B*.*napus*. However, the surface linked to petals was irregular after the petals were teared off in mutants, showing there was no abscission zone (Figure 6) (Lee et al., 2018). At the initial stage of floral organ abscission, *Bnaida* was consistent with the wild-type abscission zone cells, and after organ separation (position 17 to 24), many SECs (Secession Cells) remained on the RECs (Residuum Cells) of *Bnaida*. When comparing the scanning electron micrographs of the whole abscission area at position 23/24, it was obvious that after organ detachment, the wild-type abscission area was completely restored to a plane (Figure 6). These results suggested that *Bnaida* mutants prevented floral organ separation by affecting the normal transdifferentiation of AZ cells.

**Figure 6.**
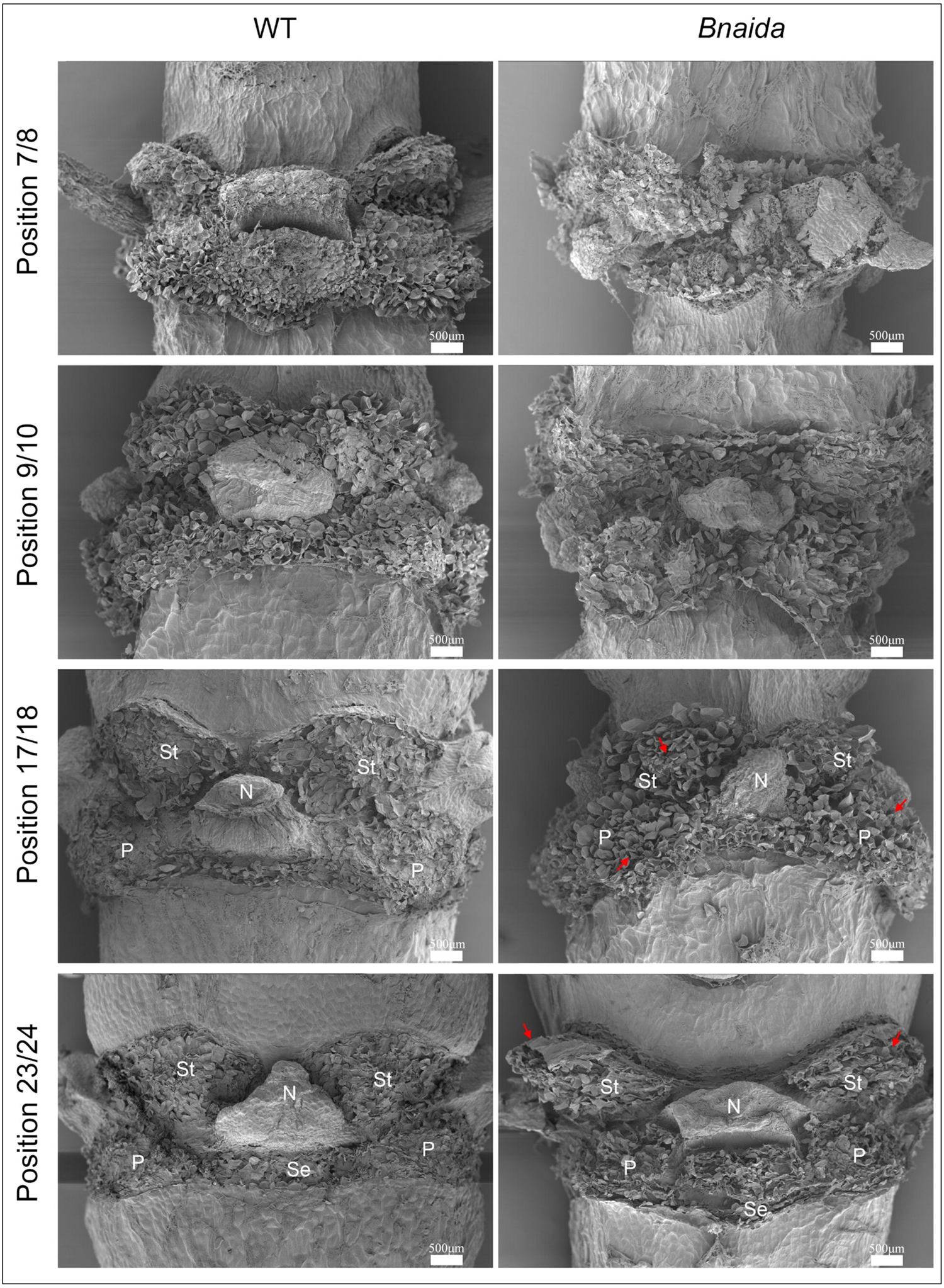
Floral AZ Scanning Electron Micrographs of *Bnaida* and WT. Scanning Electron Micrographs of Floral AZs of *Bnaida* mutants and WT. St, Stamen AZ; P, Petal AZ; Se, sepal AZ; N, Nectaries AZ. The red arrows highlight the residual floral organs. Bars = 500 μm.

### RNA-seq analysis of T2 generation mutant of *Bnaida*

To accurately analyze the difference of mRNA expression between mutants and WT, elaborating the molecular mechanism of IDA signal pathway in *B*.*napus*, large scale transcriptomic patterns of T2 mutants compared to WT lines are analyzed by RNA-seq. Clean Reads were obtained, the percentage of high-quality Clean Reads in Raw Reads in each sample was 86% to 92%; Q20 was greater than 95%, Q30 is greater than 89%, and the correct rate of base recognition was high, indicated that the sequencing results were stable and reliable. There were 77703 expression genes exceeding the threshold in WT vs *Bnaida*, of which 3778 genes were specifically expressed in *Bnaida* and 3312 genes were specifically expressed in WT (Figure S4). Subsequently, based on FDR < 0.05 and |log 2 FC| > 1 as the standard, the genes screened are differentially expressed genes (DEGs). After comparison between the two groups of samples, a total of 91 DEGs were identified, of which 49 were significantly up-regulated and 42 were significantly down regulated (Figure S5; Table S2). Then, we performed KEGG enrichment analysis on the genes specifically expressed in *Bnaida* (Figure 7a), Combined with Qvalue and the enrichment number of candidate genes, we found that the genes were mainly enriched in plant MAPK signal pathway and plant pathogen interaction. In order to further explain the expression pattern of IDA signaling pathway in *B*.*napus* and verify the reliability of RNA-Seq data, we analyzed the expression levels of related differential genes (Figure 7b). The results of qRT-PCR showed that in *Bnaida*, the expression of receptor proteins *HAE* and *HSL2, MAPK6* and transcriptional regulators *BP/KNAT1* and *KANT6* were significantly up-regulated. While, the expression of an endoglucanase gene *CEL1* related to cell wall degradation was significantly down-regulated.

**Figure 7.**
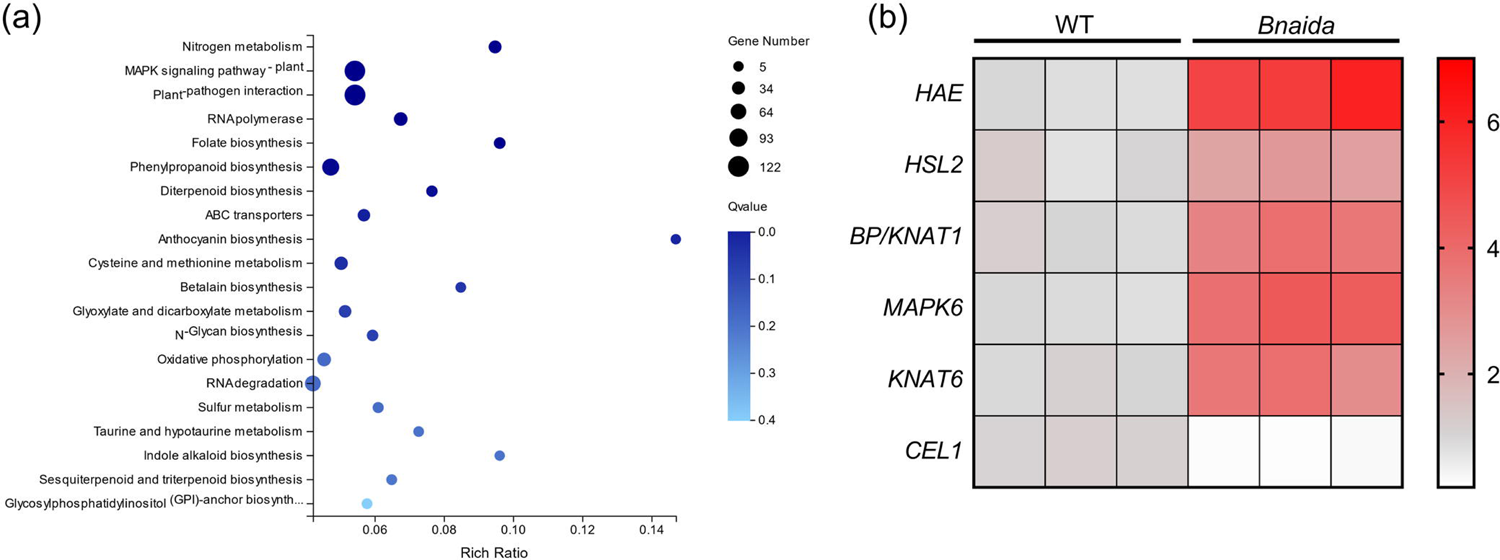
RNA-seq and differential gene expression analysis in T2 mutants. (a) 3778 genes specifically expressed in mutants were analyzed by KEGG enrichment analysis. The X-axis was enriched and the Y-axis was KEGG Pathway. The size of the bubbles indicates the number of annotated genes, and the color represents Q-value enrichment. The deeper the color, the smaller the Q-value. (b) Quantitative RT-PCR analysis of the expression of *HAE, HSL2, BP/KNAT1, MAPK6, KANT6 and CEL1* genes in WT and *Bnaida*. Color mapping represents the amount of gene expression. The gene expression of *Bnaida* shown in the figure was significantly different from that of the WT.

### Analysis of Off-target activity and main agronomic traits in *BnaIDA-A07/C06-edited* mutants

*B*.*napus* is an allotetraploid, which is likely to occur nonspecific editing. By using CRISPR-P website(http://cbi.hzau.edu.cn/cgi-bin/CRISPR) to explore the potential off-target sites in the genome were searched for all target sites, and then corresponding to the *B*.*napus* genome (http://www.genoscope.cns.fr/brassicanapus/) to obtain the matched gene sequence(K. Zhang et al., 2019; X. Zhang et al., 2021). The predicted results showed that all the Off-target sites were less than 4 SNPs compared with the *B*.*napus* genome, and all Off-scores were below 1.2%. Among the 38 predicted Off-target sites, only the Off-target on BnaC01g29010D(OT1-15) and BnaA10g08600D(OT2-3) may occurred in the CDS. Consequently, we designed primers (Table S1) and the PCR products of these two off-target sites from mutants and WT were sequenced. As expected, no Off-target genome editing in mutations (Figure S6). These results indicated that the pKSE401-BnaIDA/Cas9 system performed high-quality specific genome editing of target in *B*.*napus*.

To further investigated whether editing *BnaIDA-A07/C06* could affect *B*.*napus* growth. we compared the phenotype, 1000-seed weight, seed number per silique, branch number, plant height and germination rate between *Bnaida* and WT. The results showed that there was no significant change in the phenotype of T2 generation plants (Figure S7a). The 1000-seed weight for the *Bnaida* was 2.680-2.898g, and that for the WT was 2.682-2.888g (Figure S7b); the seed number per silique for the *Bnaida* was 12-17, and that for the WT was 12-18 (Figure S7c); the branch number for the *Bnaida* was 5-9, and that for the WT was also 5-9 (Figure S7d); and the plant height for the *Bnaida* was 92-102cm, and that for the WT was 92-100cm (Figure S7e); in addition, the germination rate for the *Bnaida* was 96-98%, and that for the WT was 96-98% (Figure S7f). Statistically, there was no significant difference in the main agronomic traits between the *Bnaida* and the WT.

## Discussion and conclusion

In this study, we successfully edited the plant small signaling peptide genes *BnaIDA-A07/C06*, which are vital for floral abscission and silique dehiscence using CRISPR/Cas9 system. We obtained stable genetic mutant lines without exogenous T-DNA. Our results showed that *Bnaida-A07/C06* mutants gained the phenotypes of *S*.*sclerotiorum* avoidance, non-floral abscission, and the silique dehiscence resistance. This is a novel germplasm resource achieved by CRISPR/Cas9 technology, with the elite agronomic traits in *B*.*napus*, such as rot stem disease avoidance mechanized harvesting friendly and sightseeing industry compatible.

There are many plants hormone signals that affect plant abscission. In *Arabidopsis*, the application of NAA will promote the early abscission of some fruits. Abscisic acid ABA indirectly affected the occurrence of abscission process by ethylene (Butenko et al., 2006; Niederhuth et al., 2013). Mutations related to ethylene perception or auxin response can lead to delayed abscission (Ellis et al., 2005). Senescence pathway is also related to plant abscission, transcription factor CDF4 promotes floral organ abscission by activating polygalacturonase *PGAZAT* gene. Driven by interconnected positive feedback circuits, CDF4, ABA and ROS promote plant leaf senescence and floral organ abscission (Xu et al., 2020). The floral organ abscission was also regulated by IDA-HAE/HSL2 pathway, the polypeptide ligand IDA binds to the LRR protein kinase HAE/HSL2 co-receptor on the cell membrane, and then phosphorylates after being recognized by SOMATIC EMBRYOGENESIS RECEPTOR KINASES(SERKs), so as to activate the downstream MITOGEN-ACTIVATED PROTEIN KINASE(MAPK) protein kinase cascade and finally control the hydrolase activity in the cell wall and regulate floral organ abscission (Meng et al., 2016; Shi et al., 2011). In this study, mutation on *BnaIDA-A07/C06* caused that attached petal and shattering proof silique in rapeseed (Figure 4a-c), we also obtained overexpression plants and found that the rate of floral organ falls off was faster than WT and mutants (data not shown). These results showed that BnaIDA was involed in the floral organ abscission. The phenotype of attached petal is a novel and elite trait that was not find in the *Brassica napus* population yet. The expression profile of *BnaIDA-A07/C06* was mainly in flower indicating it functioned within flower, subsequent KEGG enrichment analysis found that the genes were mainly enriched in plant IDA signal pathway. Editing *BnaIDA-A07/C06* in *B*.*napus* genome did not cause global change at transcriptional level. The agronomic trait investigation also confirmed that gene editing of *BnaIDA-A07/C06* had no negative effects on agronomic traits. Therefore, *BnaIDA-A07/C06* is a promising the target gene to create the novel rapeseed germplasm.

*Sclerotinia sclerotiorum* can directly infect rapeseed tissue. After the initial infection of mycelium, it will quickly spread and move inward along the petiole to the stem, resulting in water-immersion lesions, and then form a large area of mycelium mass, and finally form sclerotia in the stem, resulting in hollow stem and death of the whole plant (Bolton et al., 2006). *S*.*sclerotiorum* is one of the “cancers” of rapessd, which will directly lead to plant death, reduced production and huge economic losses. It will cause a loss of 8.4 billion yuan per year in China alone (X. Zhang et al., 2021). it is observed by scanning electron microscope that the ascospores of *S*.*sclerotiorum* can germinate and form hyphae only on the petals. If petals directly fall on the leaves of rapeseed, it cannot germinate and form hyphae and cannot form an infection pathway (I. JAMAUX et al., 1995). In the withered floral organs, the carrier rate is the highest. As the floral organs fall off, on the stems and leaves, the rapeseed will be infected and led to rot stem. Although *S*.*sclerotiorum* was also controlled through apetalous trait (without petal) to avoid the disease due to its complexity of genetics and the inferior trait linkage, the apetalous rapeseed was not popular. Hence, our study provides an novel approach to prevent *S*.*sclerotiorum* by *Bnaida* mutant.

Briefly, our work successfully utilizes CRISPR/Cas9 technology to edit in rapeseed, which greatly shortens the acquisition cycle of new germplasm. We provide novel germplasm with disease avoidance, silique shattering resistance and flowering period elongation in *Brassica napus*.

## Materials and methods

### Plant materials and growth conditions

The rapeseed cultivar Y127 was used as the transformation recipient in the present research, which was provided by Prof. Dengfeng Hong from Huazhong Agriculture University (Wuhan, China). All tissue culture was kept at 24 ± 2 °C under the light tensity of 4500 lux and light cycle of 16 h light/8 h darkness. All the transgene and WT lines were grown in the greenhouse at 24 ± 2 °C under the light tensity of 6000 lux and light cycle of 16 h light/8 h darkness.

### Fungal pathogens

Fresh *S*.*sclerotiorum* were collected from oilseed rape stems in the field in Zhenjiang, China and grown on potato dextrose agar (PDA) under 28°C.

### Vector construction and agrobacterium-mediated rapeseed transformation

In this study, the CRISPR-P 2.0 design tool (http://cbi.hzau.edu.cn/CRISPR2/) was used to design the sequence-specific targets of *BnaIDA* and to predict off-targets. The two targeting sequences driven by U6 promoters were constructed next to 35S promoter driven sgRNA in vector pKSE401(Figure 1a), which was provided by Prof. Dengfeng Hong from Huazhong Agriculture University (Wuhan, China). The genome editing of Y127 rapeseed plants by CRISPR was performed using the Agrobacterium tumefaciens-mediated(GV3101)transformation technique, according to the method previously described by Yulong Li(Li et al., 2021). The transformed plants were screened by kanamycin resistance marker in the T-DNA region of the vector pKSE401.

### Primers

Sequences of all primers used were shown in Table S1.

### RNA extraction, cDNA synthesis and quantitative real-time PCR (qRT-PCR)

Total RNA was extracted from rosette leaves and floral (position 4 to 8) using Trizol reagent. Then, cDNA was prepared by HiScript III 1st Strand cDNA Synthesis Kit (R312-02, Nanjing Vazyme Biotech, China). QRT-PCR was performed by SYBR qPCR Master mix (Q331-02, Nanjing Vazyme Biotech, China) using Applied Biosystems® QuantStudio®3 Real-Time PCR Instrument (Thermo Fisher Scientific, USA). *BnaACTIN* was used as reference gene. Relative expression was calculated automatically by the software of qRT-PCR machine based on 2^-^△△^Ct^ method.

### Identification of CRISPR-edited plants

Genomic DNA was extracted from the leaves of CRISPR transformed and WT plants according to Cetyltrimethylammonium Bromide method. Firstly U626-IDF/U629-IDR primers and Cas9-F/R primers (Supplementary Table 1) were used to identify whether it contained U6 promoter sequence and Cas9 protein sequence respectively to confirm that the transformed plants gained successful intake of CRISPR system. To further explore whether the sequences of targeted genes are modified with indels genomic fragments containing the targeted sites of *BnaIDA* were amplified by PCR with specific primers, purified with Fast pure gel DNA extraction mini kit (DC301, Nanjing Vazyme Biotech, China) and then cloned into the pMD19-T vector (TaKaRa, Japan). The CRISPR-edited sequences were screened by Sanger sequencing of pMD19-T vector.(K. Zhang et al., 2019)

### Off-target mutation analysis

The CRISPR-P design tool (http://cbi.hzau.edu.cn/cgi-bin/CRISPR) was used to predicted all potential off-target sites in T1 generation of *Bnaida* mutants. PCR products were amplified with specific primers based on the off-target sites and purified products were cloned into pMD19-T vector (TaKaRa, Japan) for Sanger sequencing. The sequences of the potential off-target sites were compared with sequences.(X. Zhang et al., 2021)

### Statistics of main agronomic traits

The mutants and WT were planted at the same time. After growing for 80 days under the condition of 16h light and 8h darkness, the plant height, number of branches and number of seeds per silique of at least 10 plants were measured. After growing for 100 days, they were harvested. After ripening for 10 days, mature seeds were obtained, and 1000-seed weight was measured. The mature seeds of mutants and WT were screened to remove the withered and stunted seeds. Lay a layer of filter paper in a 120mm glass plate and wet it. Take 50 mutant and WT seeds and arrange them in order. After 14 days of avoiding light at room temperature, count the germination rate (three replications).

### Bioinformation analysis

*BnaIDA* and its homologous sequences were obtained from the genome of *Brassica napus* ZS11 (http://cbi.hzau.edu.cn). The phylogenetic tree was constructed with MEGA 7.0 by using the Neighbor-joining method (Supplementary Figure 6).

### Measurement of forces required for silique dehiscence

The mature siliques of both WT and mutants were taken when the siliques began to turn yellow and dehydrate around 40 days after flowering. Siliques were placed under an environment with ambient temperature of 25°C and humidity of 50% for a week to keep tests on different siliques comparable prior to the measurement. Then one silique was glued to a plate, and the replum was paralleled to the plate, and the carpopodium was outside the plate. The force required for silique dehiscence was measured according to Yulong Li(Li et al., 2021). The L-shaped hook was fixed on the probe of the texture analyzer (Texture Analyzer TAXT2-HD, StableMicro System, UK). During the measurement, the plate was fixed and the L-shaped hook was moved upward at a uniform speed of 1 mm / min. When contacting the carpopodium, the L-shaped hook was moved upward at a uniform speed of 0.5 mm/min and the silique was opened. Before silique cracking, the force increases continuously, and after silique cracking, the force decreases suddenly. The peak value of the force was the maximum silique cracking force data. (Li et al., 2021; Tan XL, 2006)

### Scanning Electron Microscopy

At least 5 independent AZs of each position of flowering were collected from wildtype and mutant plants for analysis. Each position was fixed in 2.5% glutaraldehyde (w/v) in a 0.2M phosphate buffer (pH 7.2) at 4 °C then the specimens were dried using an ES-2030 freeze dryer (Hitachi) and mounted on carbon tape and sputter coated with gold palladium using an E-1010 (Hitachi). Then, samples were observed and recorded at by a S-3000N field emission scanning electron microscope (Hitachi) with a 5K accelerating voltage.

### Plant inoculation

Mycelia of *S*.*sclerotiorum* were cultured on PDA at 25°C, 7 days until black spores cover the whole plate. The Agar plug with diameter of 3mm was cut off at the edge of the growing fungi and placed upside down on the petals of flowers at the stages of position 3/4. Three independent T2 lines of CRISPR edited plants and WT were inoculated(Wang et al., 2014). The experiment was conducted according to a completely randomized block design repeated independently for 3 times. After inoculation, Brassica plants were grown in the incubator at 24 ± 2 °C with 85%-90% relative humidity and under the light intensity of 1000 lux and the light cycle of 12 h light/12 h darkness. The phenotype was observed every 24 hours after *S*.*sclerotiorum* infection.

## Ackownledgements

We thank Professor Deng-Feng Hong of Huazhong Agricultural University for providing Y127 rapeseeds and Professor Liang Guo of Huazhong Agricultural University for the pKSE401 vector.

## Author contributions

RG designed the research, performed all experiments, analyzed the data and wrote the manuscript. RS, LL, YS, YX-X, KM-Z, JW and WZ contributed to designed the research and analyzed the data. LZ-X provided the Texture Analyzer TAXT2-HD instrument. CL-S, ZW, YL-L revised the manuscript. XL-T and RB.A revised and supervised the manuscript. All authors who contributed to the study have read and approved the final version of the manuscript.

## Supplemental data

**Figure S1** FPKM values of five homologous genes (BnaA07g27400D, BnaC06g29530D, BnaC02g18450D BnaC04g26010D, BnaA02g13980D) in root, leaf, bud, silique, stamen, new pistil, blossomy petal, wilting pistil, stem, sepal, ovule and pericarp.

**Figure S2** Detection of Cas9 sequence in T1 and T2 generations. The Cas9 sequence was amplified by PCR. In the left figure, the red box indicated that plants without Cas9 were obtained in T1 generation; and in the right figure, Cas9 cannot be detected in all T2 generations. W-1/2 and WT represented wild-type plants DNA and ddH2O was used as a negative control, “+”: pKSE401-Cas9 plasmid was used as a positive control.

**Figure S3** Time-force curve record exterior forces necessary for silique dehiscence. The tensile force increased as the probe pulled the silique, and when the silique ripped, the force on the silique decreased rapidly, and the peak value records the maximum external force required for silique dehiscence.

**Figure S4** Venn diagram of T2 mutants and WT. Among the 77703 genes, 3778 were specifically expressed in mutants and 3312 in WT.

**Figure S5** The scatter plot shows that only 91 genes with significant differences among the 77703 genes, of which 49 were up-regulated and 42 were down regulated.

**Figure S6** Off-target sites analysis. All the predicated off-target sites showed less than 4 SNPs difference compared with the *B*.*napus* genome, and all off-scores were below 1.2%. Among the 38 predicted off-target sites, only the off-target on BnaC01g29010D (OT1-15) and BnaA10g08600D (OT2-3) may occurred in the CDS.

**Figure S7** Phenotype and main agronomic traits of *Bnaida*. (a) 60-day-old seedling of *Bnaida* and WT. (b) The thousand seed weight of the *Bnaida* and WT. (c) Seed numbers per silique of *Bnaida* and WT. Three individual plants of each mutant and WT were sampled, and ten siliques of each plant were counted. The values are presented as the means ± s.d for n=3 replicates. (d) Branch number of *Bnaida* and WT. Three individual plants of each mutant and WT were counted. (e) Plant height of *Bnaida* and WT. Three individual plants of each mutant and WT were measured. (f) Germination rate of *Bnaida* and WT. Three individual plants of each mutant and WT were seeded on wet filter paper, and the germination rate was counted after one week.

**Table S1** Primers.

**Table S2** Information of differentially expressed genes.

