## Supplementary figures and images for "The trifecta of disease avoidance, silique shattering resistance and flowering period elongation achieved by the *BnaIDA* editing in *Brassica napus*"

### Figure S1

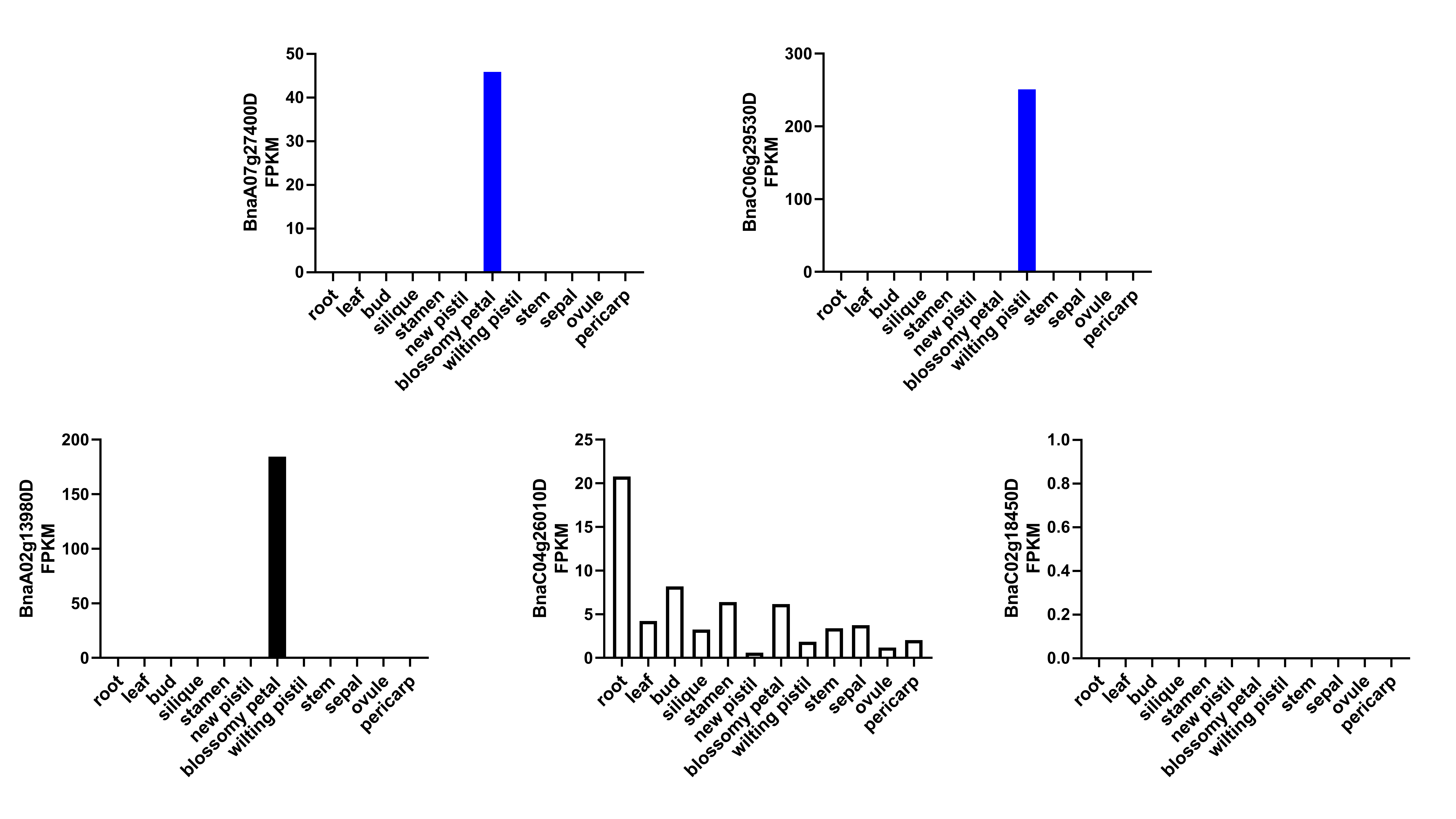

### Figure S2

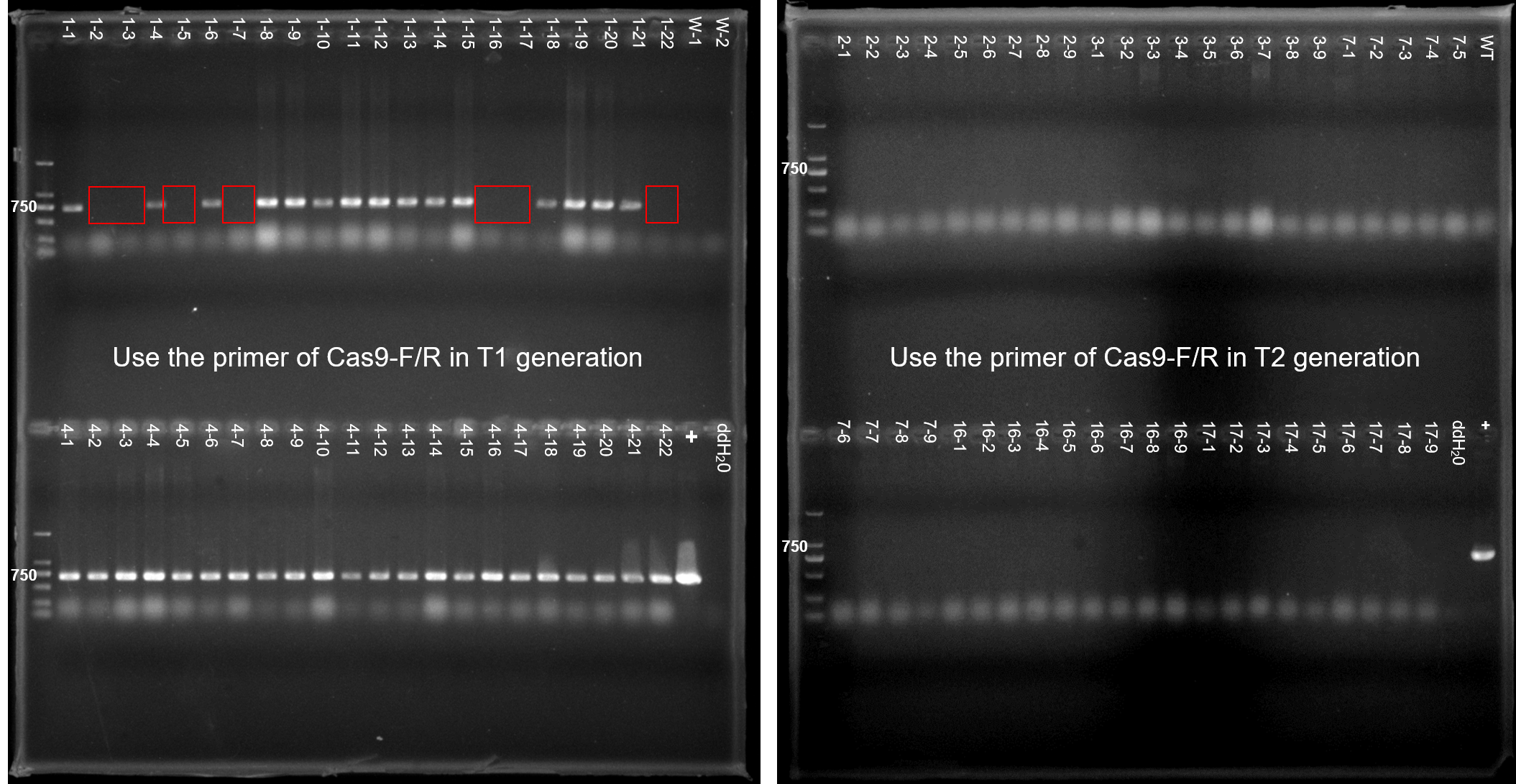

### Figure S3

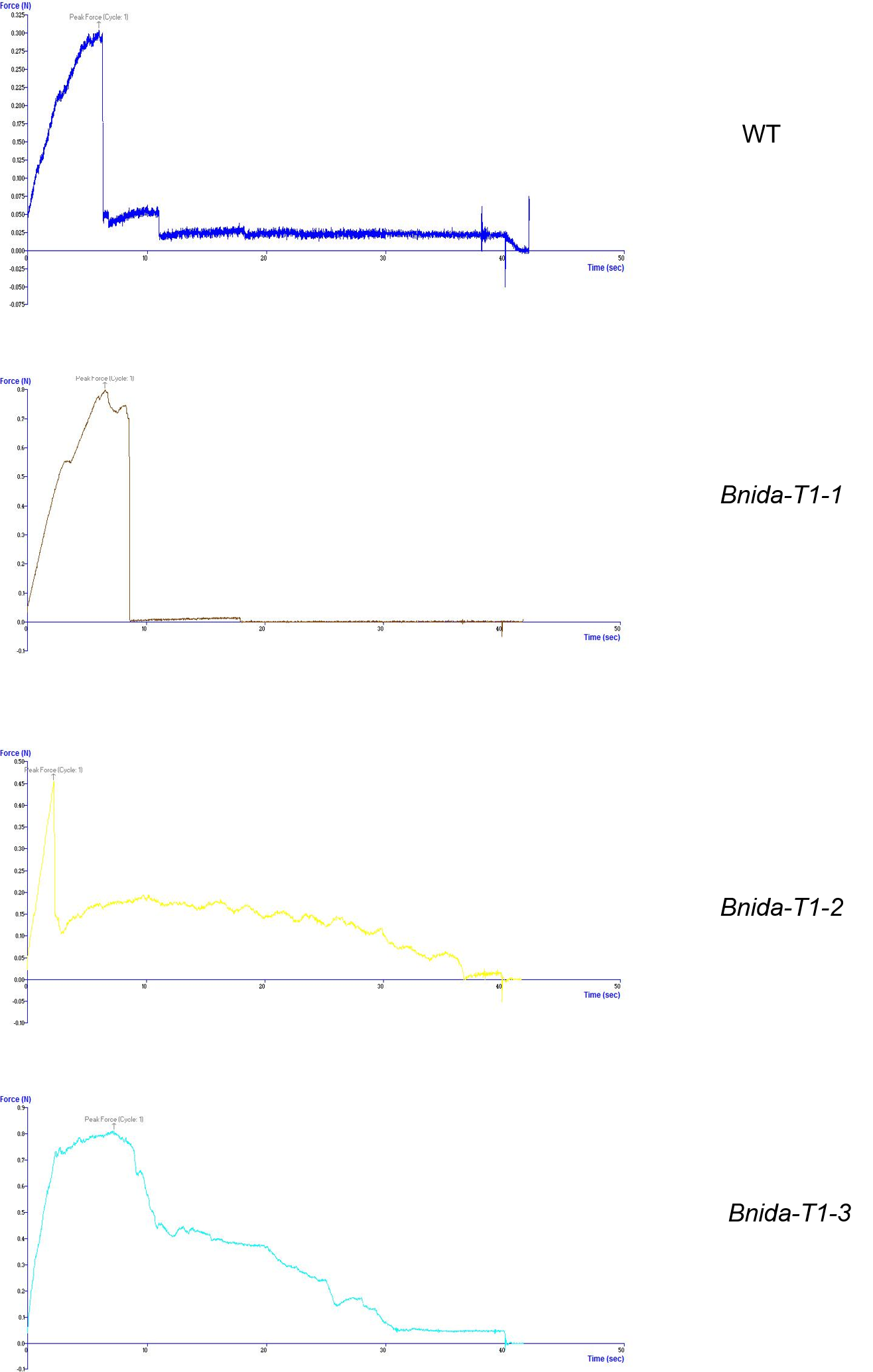

### Figure S4

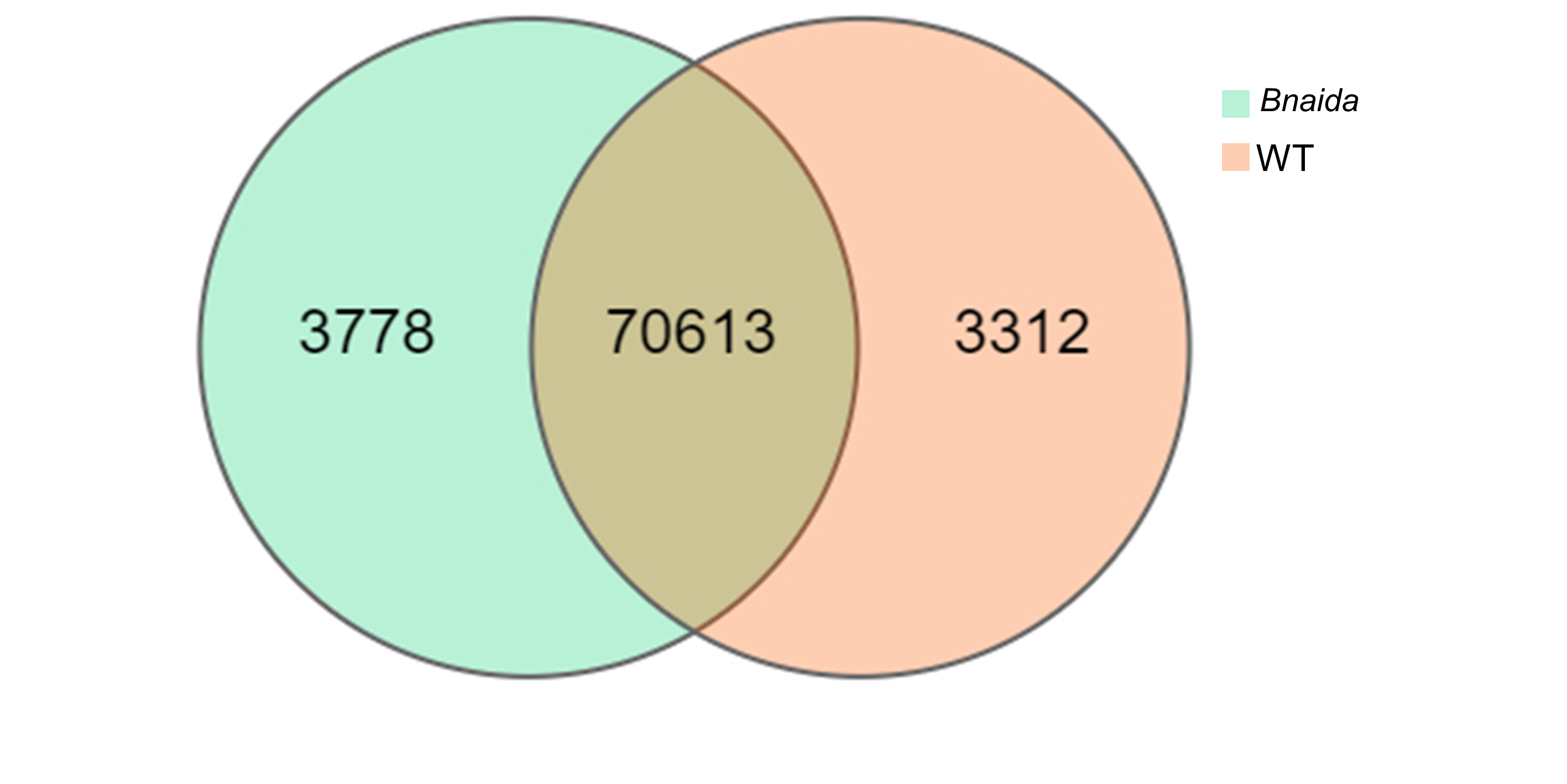

### Figure S5

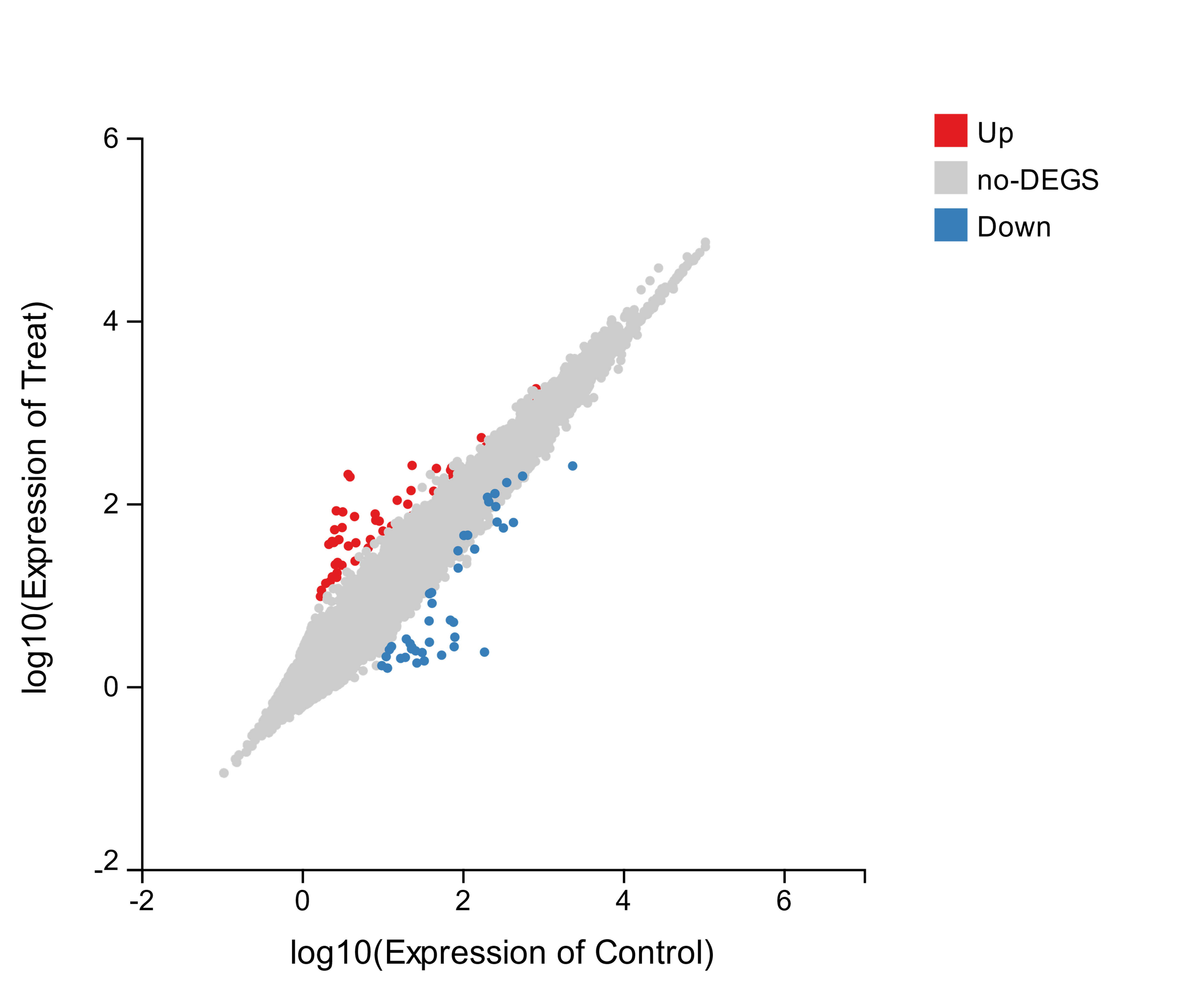

### Figure S6

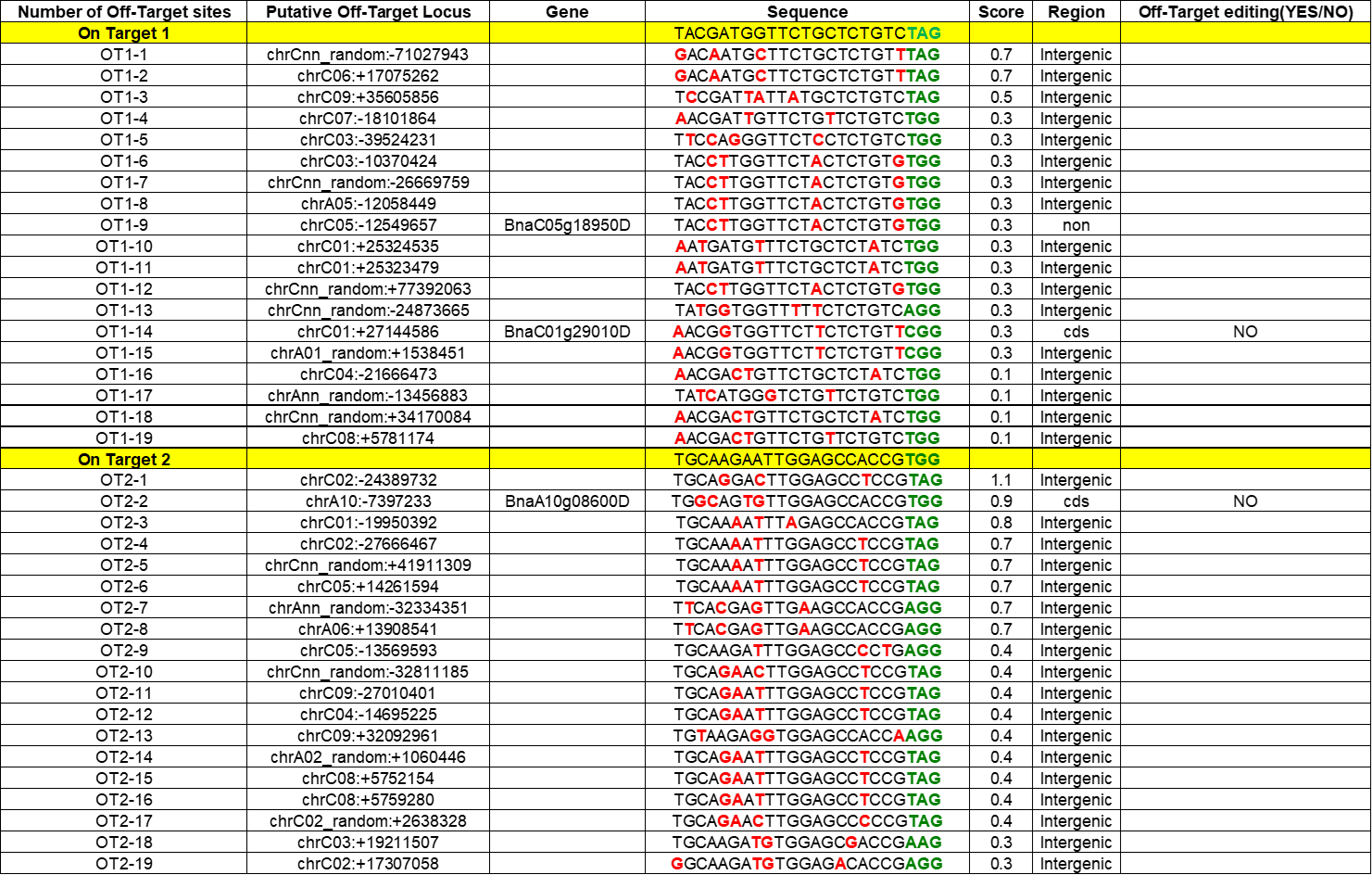

### Figure S7

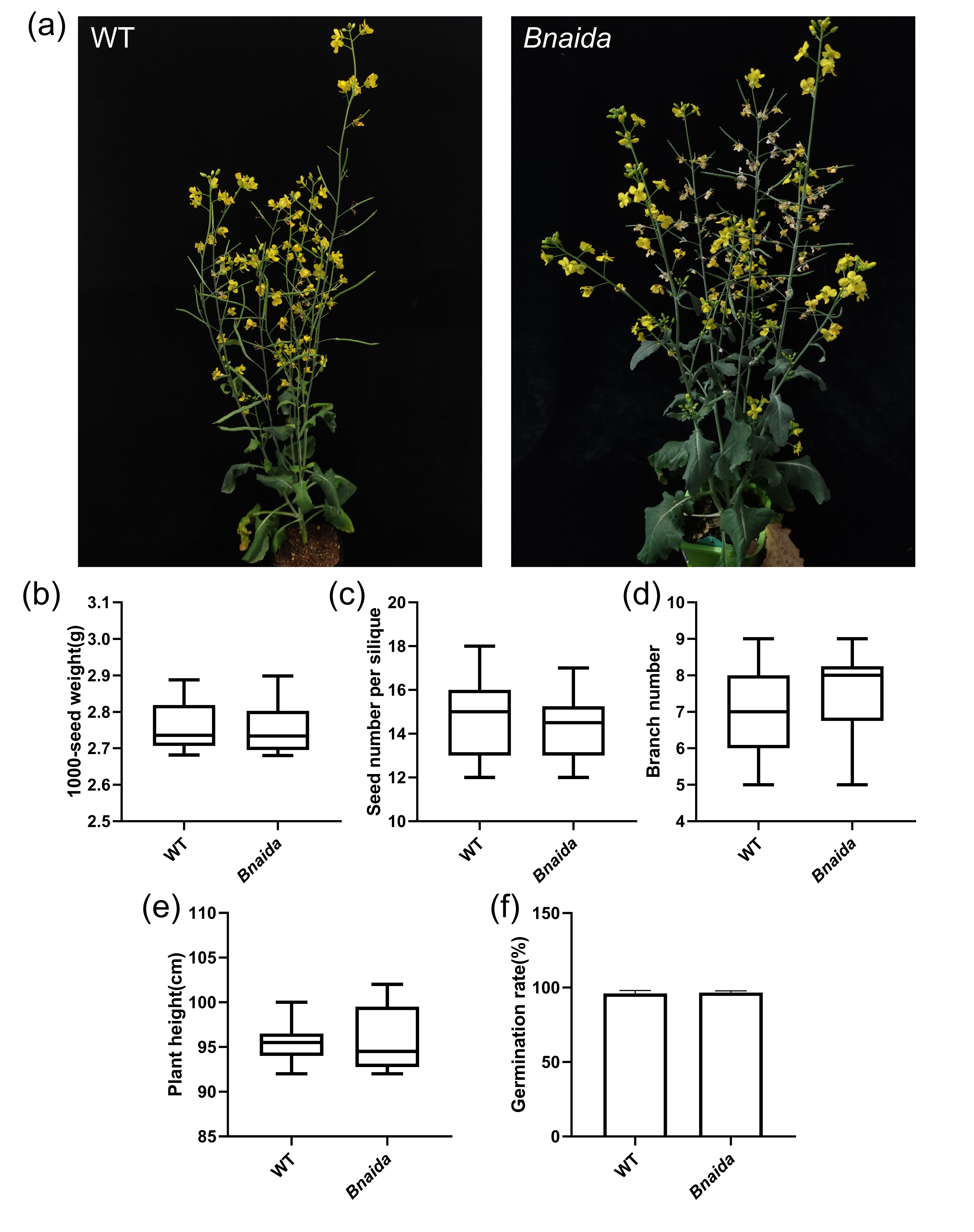
